# Sample-specific haplotype-resolved protein isoform characterization via long-read RNA-seq-based proteogenomics

**DOI:** 10.1101/2025.11.21.689818

**Authors:** David Wissel, Gloria M. Sheynkman, Mark D. Robinson

## Abstract

Protein isoform inference from bottom-up mass spectrometry (MS) relies on database search strategies that assume the reference protein database accurately reflects the full repertoire of genetic and transcriptomic states present in the sample being analyzed. Long-read RNA sequencing (lrRNA-seq) now enables simultaneous recovery of complete transcript (splice) structures and the genetic variants present on each molecule, offering a direct route to allele-specific isoforms. Yet, this capability has not been fully leveraged to improve MS-based proteogenomics workflows. Here, we develop an end-to-end workflow for constructing and searching haplotype-resolved, sample-specific proteomes using matched lrRNA-seq and MS data. We benchmark phasing algorithms on PacBio lrRNA-seq from Genome-in-a-Bottle samples and identify methods that achieve high phasing accuracy and completeness. Our Snakemake pipeline leverages existing methods to perform variant calling, read-based phasing, transcript discovery, haplotype-resolved proteome construction, MS search, and downstream annotation. To demonstrate its utility, we apply the workflow to an induced pluripotent stem cell line (WTC11) and to an osteoblast differentiation time course. We show that sample-specific haplotype-resolved databases enable the detection of variant and splice peptides, allele-specific protein isoforms, and linked variants not detectable with reference proteomes. Together, our results demonstrate that lrRNA-seq-based phasing is feasible and effective for proteogenomics and provide a practical framework for allele-resolved proteome characterization in dynamic or disease-relevant settings.

## Background

The accurate identification and characterization of proteoforms has been a core focus of the proteomics and proteogenomics field [1]. Although top-down mass spectrometry (MS) methods have become increasingly popular for the direct detection of proteoforms [2], the global characterization of protein isoforms still primarily relies on bottom-up, or shotgun sequencing, regimes in which the existence of proteoforms can only be inferred from identified peptide fragments [3, 4]. In such approaches, the short peptide fragments are mapped back to the annotated sequences in a reference proteome, and the protein sequences with the highest support, based on the inference algorithm, are reported. The accuracy of such inference methods depend on the extent to which the reference proteome reflects the actual composition of the proteome in the sample subjected to analysis. However, reference proteomes are incomplete; the number of proteoforms in the human proteome is estimated to be in the hundreds of thousands, if not millions, owing primarily to genetic variants, alternative splicing, and post-translational modifications [5]. Consequently, accurate protein inference requires protein annotations that faithfully capture genetic and transcriptomic complexity in a cell or tissue specimen [4].

In human cells, the protein inference challenge is compounded by the fact that primary protein sequences (N to C-terminal AA chains) contain different coding (e.g., missense) genetic variants across individuals. Variant calling attempts to identify such variants by looking for differences to a reference genome of the same species. Genetic variation affects protein sequences not only through single-event amino-acid changes but also insertion and deletion events (indels) of polypeptide segments. Furthermore, combinations of these variants can co-occur and be co-inherited on the same chromosome. These co-inherited variants define allele-specific protein sequences, or protein haplotypes [6]. Variant phasing methods aim to reconstruct these haplotypes by determining variants that reside together on the same chromosome.

Despite their biological relevance, allele-specific protein sequences, or haplotypes, are rarely represented in standard proteome databases, which typically assume a single reference sequence per isoform. Beyond genetic variation, transcriptomic regulation adds an additional layer of complexity since a given haplotype can give rise to multiple alternatively spliced transcripts. As a result, the final amino acid sequence of a protein is jointly determined by both the set of variants present on each allele and the specific transcript (splice) structure through which those variants are expressed.

Each allele may give rise to multiple splice isoforms, meaning that the same set of variants may be incorporated, excluded, or combined in distinct ways across known and novel transcripts. An accurate protein reference for MS-based inference, therefore, requires both knowledge of the genetic variants present on each allele and the correct set of transcript structures expressed in that sample, so that the resulting protein models faithfully represent the allele-resolved sequences the cells in question actually produce. These considerations motivate the need for a systematic approach that identifies genetic variants, determines their phase, and establishes the full-length transcripts within which the genetic variants are expressed.

Phasing can be performed using several strategies, including population-based phasing by leveraging large reference panels, trio-based phasing by using parental genotypes, and read-based phasing by inferring haplotypes directly from sequencing data [7]. Of these, read-based phasing most easily enables sample-specific reconstruction of allele structure because it uses the long or linked reads obtained from the sample itself to determine which variants co-occur with eachhother.

We now give a short overview of recent proteogenomics approaches that enable the creation of sample-specific proteomes, in addition to variant phasing. Related work has developed along two largely independent axes: sample-specific proteogenomics workflows and protein haplotype prediction from genetic analysis of population cohorts. We begin with proteogenomic frameworks that refine the MS search space by constructing customized proteomes for a given sample.

Cesnik et al. [8] develop a consolidated pipeline, Spritz, that integrates short-read RNA-seq with variant calling methods to produce protein databases containing both variant and splice-derived protein isoforms. They used GATK [9] for variant calling and StringTie2 [10] for novel transcript reconstruction, and SnpEff [11] to project the variants onto protein sequences to generate proteins. Spritz enables the identification of both variant peptides and, in some cases, proteoforms by utilizing its sample-specific database in bottom-up and top-down MS searches, respectively. Spritz processes short-read RNA-seq data; thus, it is limited to incomplete transcript reconstructions that may not capture full-length isoforms. In this regard, Miller et al.

[4] introduced a pipeline for long-read RNA-seq (lrRNA-seq) data that constructs sample-specific, full-length proteomes. The authors used the Iso-Seq workflow [12] to assemble a sample-specific transcriptome, annotated open reading frames (ORFs) using CPAT [13], and construct the corresponding personalized proteome. Miller et al. [4] shows that such databases can enhance characterization of specific protein isoforms, with detailed isoform classification automatically performed based on an extension of SQANTI3, SQANTIProtein [14]. However, though capturing splicing complexity at full-length resolution, the pipeline omits genetic variation. Salz et al. [15] introduce a proteogenomic pipeline that does include genetic variants in full-length isoform models, reporting on a constructed personalized proteome for the HG001 Genome in a Bottle (GIAB) cell line [16]. Phased genetic variants were obtained from a previously published variant phasing dataset generated using an ancestry-based genetic phasing approach [17] and novel transcript structures obtained from an analysis of Oxford Nanopore data on the same cell line [18]. However, while genetic variants are incorporated into protein sequences, the framework focuses on the comparison of *de novo* and reference-based approaches and does not offer or evaluate the option to call and phase variants based on sample-specific lrRNA-seq data. Beyond these proteogenomic efforts, a separate family of strategies has focused not on sample-specific database construction, but on defining the theoretical space of protein haplotypes arising from genetic variation across human populations. Haplosaurus utilizes Variant Effect Predictor (VEP) [19] to map all variants from the 1000 Genomes dataset [20] onto translated coding sequence (CDS)s (i.e., reference proteins) [6]. This framework defines all possible protein sequences encoded by phased alleles and highlights how common haplotypes can differ substantially from canonical reference proteins across populations and lead to changes in amino acids that affect protein functions. Vašíček et al. [21] leveraged Haplosaurus to generate databases for MS searching. They generated haplotype protein FASTAs and performed MS database searches to assess the downstream detectability of peptide-level variants, identifying *≈* 176 thousand peptides at 1% FDR, including *≈* 7.5 thousand peptides containing one variant and 352 with multiple variants. This work is an early demonstration that multi-variant peptides from protein haplotypes can be observed in a systematic manner. The same authors built on this work by introducing a scalable bioinformatic pipeline, ProHap [22] and companion visualization tool [23], which constructs personalized proteomes by overlaying phased variants from population reference panels onto all Ensembl transcript annotations. The authors illustrate the usage of their method on the 1000 Genomes dataset, revealing cross-population proteome complexity. Collectively, all these studies establish the theoretical landscape of protein haplotypes and demonstrate their detectability in MS data. However, because they rely exclusively on reference transcript annotations, they cannot represent sample-specific or novel transcripts and cannot jointly model the allele-specific combinations of variants and splice choices that give rise to the proteoforms present in an actual sample.

With the increasing accessibility of highly accurate lrRNA-seq datasets (e.g., PacBio Iso-Seq, ONT cDNA), we wanted to investigate the potential to build sample-specific, allele-resolved personalized proteomes directly from experimental lrRNA-seq sequence data. In lrRNA-seq, each read corresponds to a single transcript and encodes both the complete splicing structure of the expressed isoform and the genetic variants present in that molecule. In principle, this enables the simultaneous recovery of isoform identity and variant phase from the same reads, allowing direct translation into allele-specific protein isoforms. In parallel with advances in long-read sequencing, methods for phasing genetic variants have matured [24–27], and it is now possible to employ phasing on long-read transcript data [28, 29], including methods that leverage allele specific expression (ASE) [30].

In this work, we present an end-to-end framework for constructing haplotype-resolved, sample-specific proteomes directly from long-read RNA sequence data. We briefly summarize our main contributions. First, in order to determine the ideal method for variant calling, we perform a benchmark of methods for variant phasing on two PacBio lrRNA-seq datasets, focusing on raw phasing performance (given the lack of diverse large benchmark datasets with matched MS and lrRNA-seq). Second, we implement a Snakemake workflow that enables the creation and search of haplotype-resolved, sample-specific proteomes from sample-matched lrRNA-seq and MS data [31]. Third, we characterize the proteome complexity observed in an lrRNA-seq sample from the WTC11 cell line as well as in a model of differentiation from iPSCs to osteoblasts, showing potential applications of our workflow in disease or other more complicated experimental designs. Finally, we highlight new aspects of how haplotype-resolved proteomes relate to protein isoform characterization.

## Results

### Read-based phasing methods exhibit good performance on PacBio long-read RNA-seq data

We evaluated the performance of existing methods for variant phasing using PacBio lrRNA-seq on two GIAB datasets, HG002 and HG005. Our goal was to assess which methods most accurately reconstruct haplotypes for known, ground-truth GIAB variants when applied specifically to PacBio lrRNA-seq data. Phasing accuracy was quantified using the switch error rate, which measures how often the inferred haplotype assignment incorrectly switches between parental alleles when traversing consecutive heterozygous variants along a transcript (see **Methods**). Lower switch error rates indicate more consistent haplotype reconstruction. In addition, we evaluated phasing completeness, using both the total number of phased variants and the proportion of fully phased CDSs (i.e., CDSs that consisted of a single contiguous phase block). When using lrRNA-seq data, the length of phase blocks of interest is that within the length of protein-coding genes. For downstream MS database construction, methods that phase a large number of variants but do not fully phase a large proportion of CDSs are less useful.

Overall, we found similar performance on both the HG002 and HG005 datasets. WhatsHap, HapCUT2, and Margin all performed well at low switch error rates and performed broadly similar to each other, with WhatsHap achieving a slightly worse switch error rate compared to the other two, while phasing notably more variants overall, as well as phasing a larger percentage of CDSs fully (Fig. 1A). phASER did not perform well, underperforming the top three methods based on phasing accuracy, with similar performance to WhatsHap based on phasing completeness. HiPhase performed worse than all other methods regarding phasing accuracy on HG002 and HG005, but never fully phased less than 80% of all CDSs. There were no notable differences in performance when including or excluding indels for HG002 and HG005 (Fig. 1A).

**Fig. 1.**
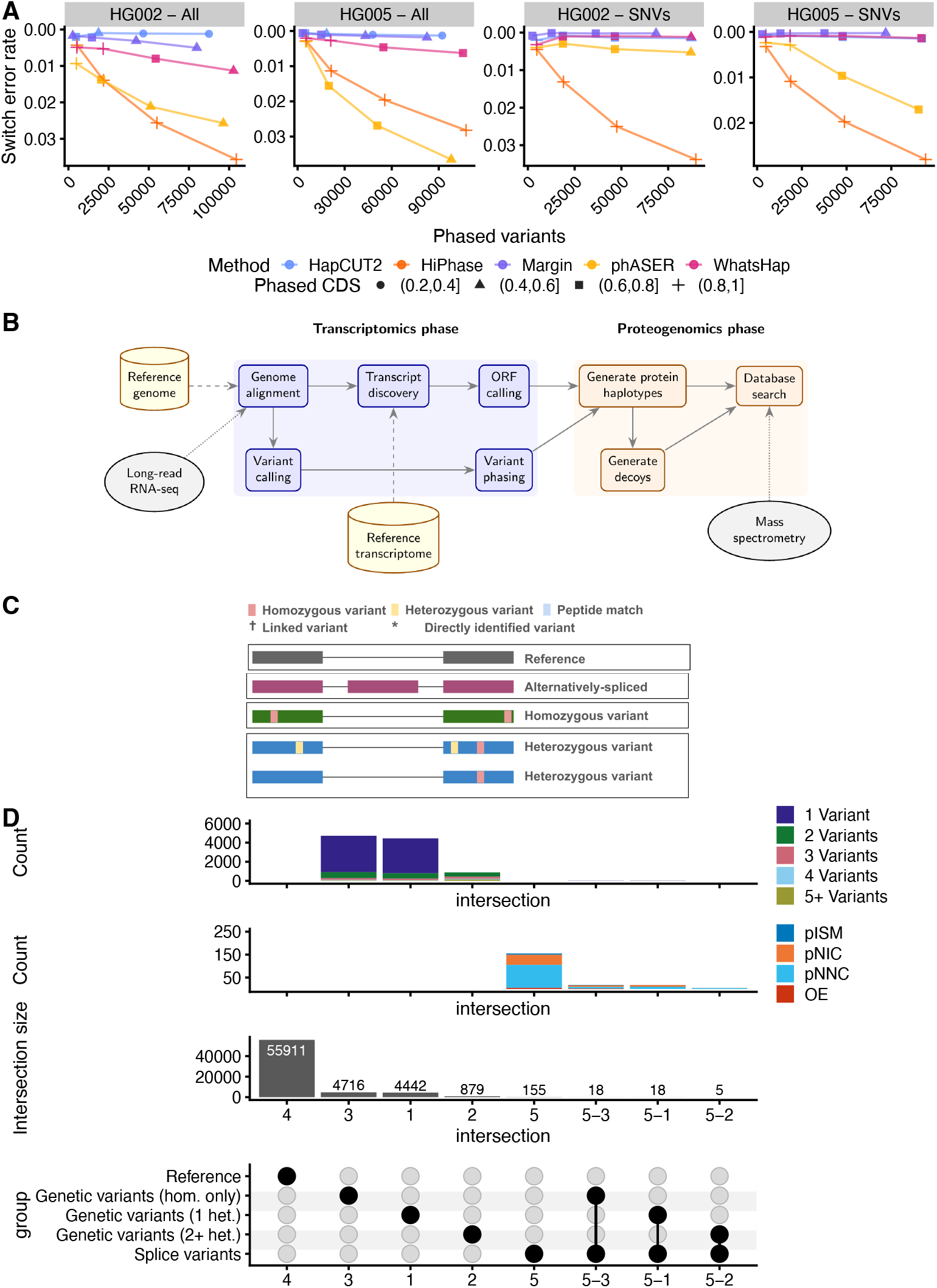
Generating sample-specific proteomes using our workflow enables a comprehensive characterization of the theoretical proteome of a system, in our case, the WTC11 cell line. **A. Phasing performance as measured by switch error rate of established phasing methods on lrRNA-seq data by the number of phased variants and fully phased CDSs, stratified by dataset and inclusion of indels. B.** Schematic view of our pipeline to perform sample-specific haplotype-resolved proteogenomics. **C**. Overview of categories of protein isoforms identifiable with our workflow. Legend also used in Fig. 4. **D**. Number of protein isoforms belonging to each variant category and the reference, along with a breakdown of complexity within each variant category.

Based on these results, for variant phasing, in particular with the goal of performing MS search with PacBio lrRNA-seq data, we recommend the usage of WhatsHap, Margin, or HapCUT2 and use WhatsHap throughout our manuscript.

### A modular workflow enables sample-specific haplotype-resolved proteogenomics

We implemented our method as a Snakemake pipeline, leveraging existing methods wherever possible and augmenting them with custom scripts and tools when necessary.

Users provide a reference genome and transcriptome, together with sample-matched lrRNA-seq and MS data (Fig. 1B). The pipeline begins by aligning lrRNA-seq data to the genome, followed by variant calling and transcript discovery, which are performed in parallel to capture genetic variation and alternative splicing, respectively. Variants are filtered based on coverage and call quality, phased using WhatsHap, and post-processed to ensure that regions of interest—typically CDSs—form contiguous phase blocks. Novel transcripts undergo an additional ORF calling step using ORFanage, followed by filtering to retain CDSs of sufficient length.

Using the resulting phased variants and CDS annotations, we next construct protein haplotypes using Haplosaurus and generate decoys for downstream MS search (Fig. 1B). First, reference protein haplotypes are generated from GENCODE annotations using gffread. Then, for the sample-specific proteome, we replace any protein isoform that has non-silent variants with the corresponding Haplosaurus-predicted protein haplotype. Lastly, we add the protein haplotypes corresponding to novel protein isoforms (whether they contain genetic variation or not).

Finally, the sample-specific database is searched against the matched MS data (Fig. 1B). Following the search, our pipeline performs post-search annotation of all protein isoforms, creates files to enable the creation of browser tracks, and performs protein inference.

### Sample-specific haplotype-resolved proteogenomics enables the identification of a broad class of protein isoforms

In contrast to competing approaches, which either rely on population-based variant data and thus might not necessarily reflect sample-specific variation or do not jointly integrate splice with phased genetic variation, our approach consolidates a broad class of proteoform candidates. Specifically, our pipeline leverages reference protein isoforms (from a reference database, such as GENCODE) and alternatively spliced protein isoforms (from transcript discovery on the inputted lrRNA-seq data). In addition, our pipeline includes protein isoforms that contain only homozygous variants and ones that contain heterozygous variants via variant phasing (Fig. 1C).

### Composition of the haplotype-resolved proteome in WTC11

We applied our pipeline to a WTC11 cell line sample. Below and throughout the remainder of the text, we only consider non-silent genetic variation, due to the centering of our work on the proteome.

As expected, the majority of protein isoforms in our WTC11 database were identical to isoforms found in the GENCODE reference database from which our database was derived (55,911, 84.5%) (Fig. 1D). Further complexity in our sample was represented primarily by genetic variants, with only 196 (0.3%) protein isoforms exhibiting alternative splicing (Fig. 1D), relative to the reference GENCODE proteome, while 10,078 (15.2%) protein isoforms presented one or more non-silent variations (Fig. 1D). In our sample, the majority of variant protein isoforms had either no heterozygous variants (4,734 or 47.0% of variant protein isoforms) or one heterozygous variant (4,460 or 44.3% of variant protein isoforms), with only a few presenting two or more heterozygous variants (884 or 8.8% of variant protein isoforms) (Fig. 1D). There was only moderate overlap between reference variation due to alternative splicing and genetic variation. In particular, a total of 41 protein isoforms (0.1%) exhibited both alternative splicing and some form of genetic variation (Fig. 1D).

Within protein isoforms harboring genetic variants, most tended to have only one or two variants, when accounting for both homozygous and heterozygous variants (Fig. 1D). In contrast, protein isoforms arising from alternative splicing were characterized based on their splice structure rather than variant count. Under this classification, the majority of splice variants being categorized as protein novel-not-in-catalog (pNNC), followed by novel-in-catalog (pNIC), incomplete-splice-mismatches (pISM), and orphaned exons (OE) (Fig. 1D) when categorizing their protein isoform structure.

### Long-read RNA-seq-based personalized proteomes enable improved protein isoform identification and characterization

In MS searches of our sample-specific haplotype-resolved database, as well as reference UniProt and GENCODE databases (see Methods), we found that all databases were largely concordant in terms of identified peptides (Fig. 2A). In particular, 122,951 identified peptides were common to all databases, representing 98.8%, 95.8% and 99.4% of sample-specific, UniProt, and ENCODE peptides, respectively. Overall, the UNIPROT database identified the most peptides, totaling 128,336, while the sample-specific database and GENCODE identified 3,906 and 4,682 fewer, respectively. Peptides that were detected (1% FDR threshold) in only one database were often completely absent from other databases, especially UNIPROT, which had many peptides that were absent from both GENCODE and the sample-specific database (Fig. S1A-B). In contrast, those peptides detected in two databases were typically present in the remaining database but narrowly missed the FDR cutoff of 1%. Overall, in the search against the sample-specific database, we found peptide-level FDRs to be well calibrated in separating between decoys and non-decoy protein isoforms, both overall and stratified by peptide categories (reference, alternative splicing-peptides, and variant-associated) (Fig. S1C).

**Fig. 2.**
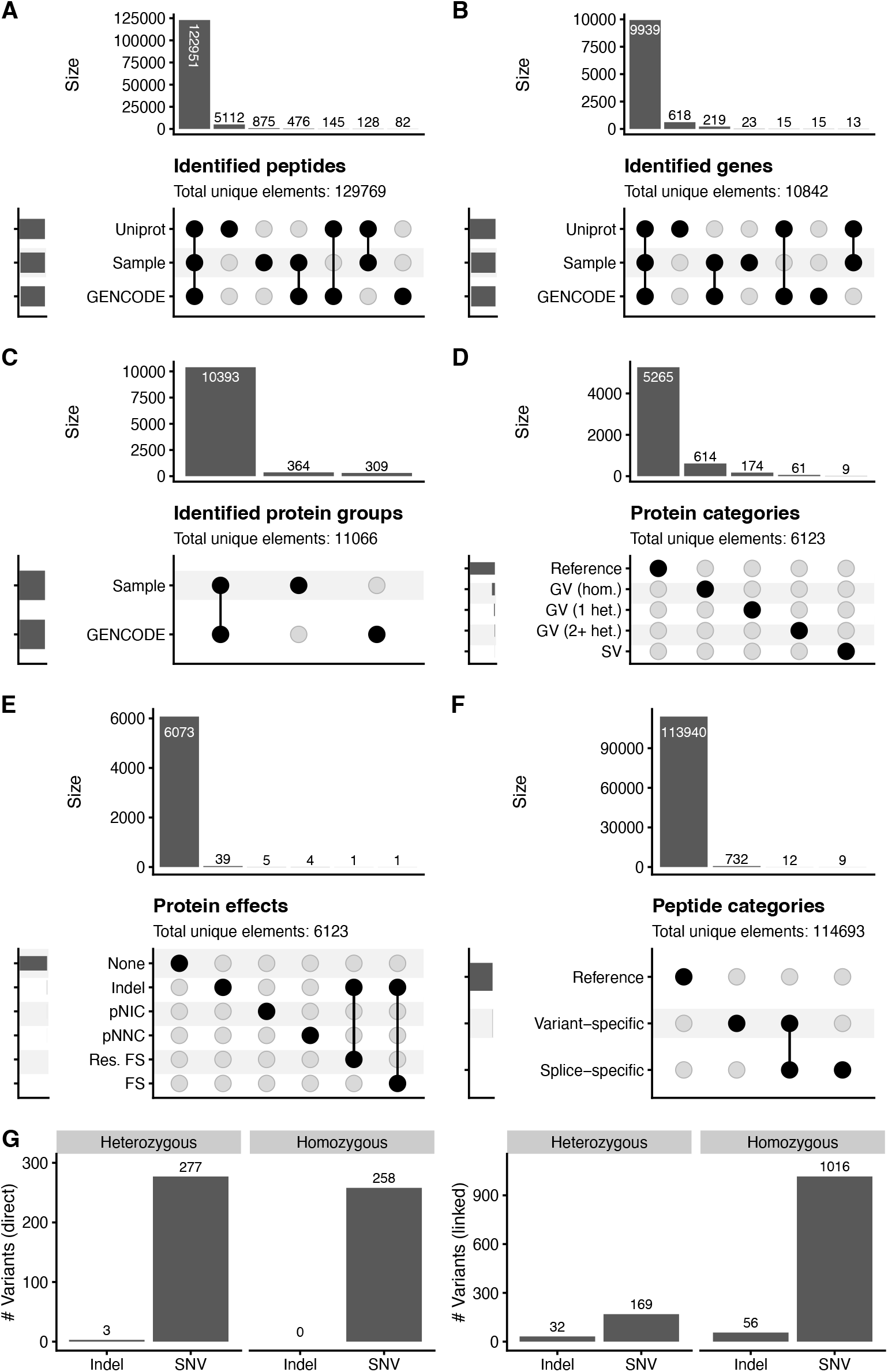
Searching MS data against the sample-specific proteome of the WTC11 cell line reveals good concordance with reference proteomes, low linkage between protein isoform types and protein effects, and identification of both direct and linked variants. **A.** UpSet plot of peptide overlaps between sample-specific proteome, UniProt, and GENCODE reference. **B**. UpSet plot of gene overlaps between the sample-specific proteome, UniProt, and GENCODE reference. **C**. UpSet plot of protein group overlaps between the sample-specific proteome and GENCODE reference. **D**. UpSet plot of protein isoform types between alternative splicing and variant isoforms. **E**. UpSet plot of overlaps of protein effects caused by genetic variation and alternative splicing. **F**. UpSet plot of peptide categories identified in searching the sample-specific proteome (we note that this number does not account for PTMs and is thus slightly lower). **G**. Left: Number of variants identified via direct peptide evidence, stratified by zygosity. Right: Number of variants identified via linked peptide evidence, stratified by zygosity. We counted a peptide as identified if it had a q-value of less than 0.01 and was not a decoy or contaminant. Identified genes were not compared to UNIPROT due to non-trivial gene mapping for UNIPROT. **D** and **E** are based on uniquely identified protein isoforms.

To determine whether peptide-level differences translated into differences in gene identification, we next performed gene-level analysis. We determined that a gene was identified if a protein group was identified for which all proteins were products of this particular gene. Gene-level search results largely recapitulated our peptide-level search results. A total of 9,939 genes were identified in common between all three databases (sample-specific, GENCODE, and UniProt), with UniProt identifying the most genes with 10,585, and the sample-specific database and GENCODE trailing it slightly with 10,194 and 10,188 identified genes, respectively (Fig. 2B).

Protein inference was compared only between our sample-specific database and GENCODE, as both use a consistent transcript-based protein accession, enabling direct comparison between protein groups. Protein inference identified 10,393 protein groups that were identical (that is, the protein groups contained exactly the same transcript identifiers) between the sample-specific search and the GENCODE search, with 364 protein groups being unique to the sample-specific search and 309 to GENCODE (Fig. 2C). Thus, the sample-specific search identified a total of 10,757 protein groups, 55 more than GENCODE.

To identify the source of differences between searches, we examined protein groups that were unique to one database. In most cases, protein groups unique to the sample-specific search contained at least one protein sequence that is not present in GENCODE, and therefore could not be matched to any GENCODE protein group (Fig. S1D). Differences attributable to alternative grouping of the same set of proteins—such as one group being a strict subset or superset of a group in the other database—were rare (Fig. S1D).

We then examined whether different types of sequence alterations co-occurred within the same protein isoform by restricting analysis to uniquely identified protein isoforms (that is, protein groups that only contained a single transcript id). Across these isoforms, we found no association between isoform categories (Fig. 2D) and only limited co-occurrence of distinct variant effects (Fig. 2E). Specifically, only one protein group contained both an indel and a frameshift, and one protein group presented an indel with a resolved frameshift.

Next, we examined peptide-level identifications to characterize how reference, variant, and splice-derived sequences contribute to the observed proteome. Peptides that did not match the reference sequence were classified as variant peptides, splice peptides, or both (see Methods). In particular, of the 114,421 annotatable total peptides identified with the sample-specific database, 113,940 (99.3%) were classified as reference peptides, with 744 and 21 falling under the splice-specific and variant umbrella, under our definition, respectively (Fig. 2F).

In addition to performing analyses on the protein isoform level, we also investigated the degree to which the MS search identified particular variants. We first focused on directly identified variants, wherein a unique peptide spans the variant site and directly confirms the altered amino acid sequence. The search of our sample-specific database identified a total of 538 variants, of which 258 were homozygous and the rest heterozygous (Fig. 2G, left). Furthermore, the direct identification of indels was rare, with only three heterozygous and no homozygous indel having been identified via direct peptide evidence in our search.

Beyond variants directly supported by peptide evidence, haplotype phasing enables inference of additional variants through linkage. Specifically, variants that reside on the same phased haplotype as a directly identified variant can be inferred even when no peptide spans the variant site itself. We found that variants identified by linkage were more abundant overall, with 201 linked heterozygous variants being supported in our search, of which 32 were indels and 169 SNVs (Fig. 2H). Overall, variant identification through linkage substantially expanded variant coverage, with a total of 1,072 homozygous variants having been identified, of which 56 were indels and 1016 SNVs.

### Haplotype-resolved proteogenomics enables the characterization of broad classes of protein isoforms

Overall, we show that haplotype-resolved proteogenomics approaches based on lrRNA-seq enable the characterization of broad classes of protein isoforms, including reference isoforms, alternatively-spliced isoforms, variant isoforms containing homozygous genetic variation, and variant isoforms containing one or more heterozygous variants (Fig. 3A).

**Fig. 3.**
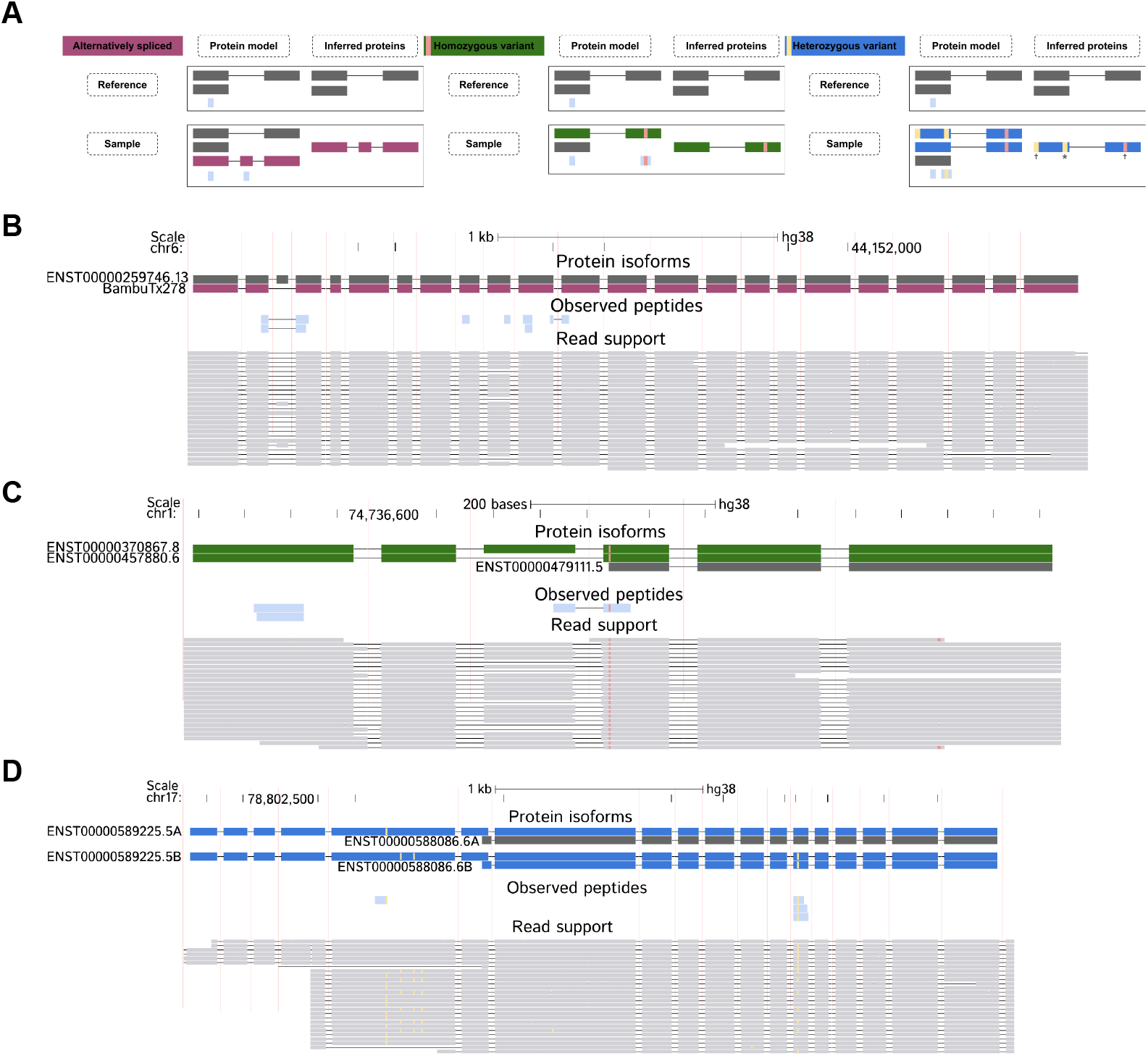
Sample-specific haplotype-resolved MS database searches enable identification of diverse protein isoform classes. **A.** Examples of protein models and corresponding inferred proteins of protein isoforms categories identifiable with our workflow. **B**. Example browser track for an alternatively spliced protein isoform on the *TMEM63B* gene. **C**. Example browser track for a homozygous variant protein isoform on the *TYW3* gene. **D**. Example browser track for a heterozygous variant protein isoform on the *USP36* gene.

Accounting for sample-specific alternative splicing events allows protein models to better reflect the set of isoforms expressed in a given sample, by excluding reference protein isoforms not supported at the RNA level, while at the same time including novel protein isoforms not present in the reference [4, 32, 33] (Fig. 3A, B). Including genetic variation further increases sequence specificity and introduces variant peptides that can contribute to protein inference [34, 35], consistent with prior proteogenomics studies (Fig. 3A, C). Lastly, resolving heterozygous genetic variation enables the construction of allele-specific protein models that could support the identification of protein haplotypes (Fig. 3A, D). Phasing additionally enables the identification of linked variants (Fig. 3A), i.e., variants inferred through co-occurrence on the same haplotype despite lacking direct peptide evidence.

### Sample-specific proteogenomics enables the analysis of condition-effect or differentiation models

Lastly, we present a vignette illustrating how our framework can be applied to multi-condition or differentiation settings. We analyzed publicly available, sample-matched PacBio lrRNA-seq and TMT-labeled MS data from an iPSC into osteoblast differentiation [36].

As an example, we highlight the ENST00000379802.8 transcript, belonging to the *DSP* gene. Consistent with the known role of DSP in cell-cell adhesion [37], we find that ENST00000379802.8 increases expression and TMT reporter ion intensities in iPSCs (Day 0) relative to osteoblasts (Fig. 4A). In addition, both datasets support the presence of both haplotypes for the transcript, with the second (B) haplotype consistently observed at a lower abundance. The two haplotypes both carry a single coding heterozygous SNV, and peptide evidence confirms the presence of both allelic protein products at the same sequence position, with and without the heterozygous variant in question (Fig. 4B).

**Fig. 4.**
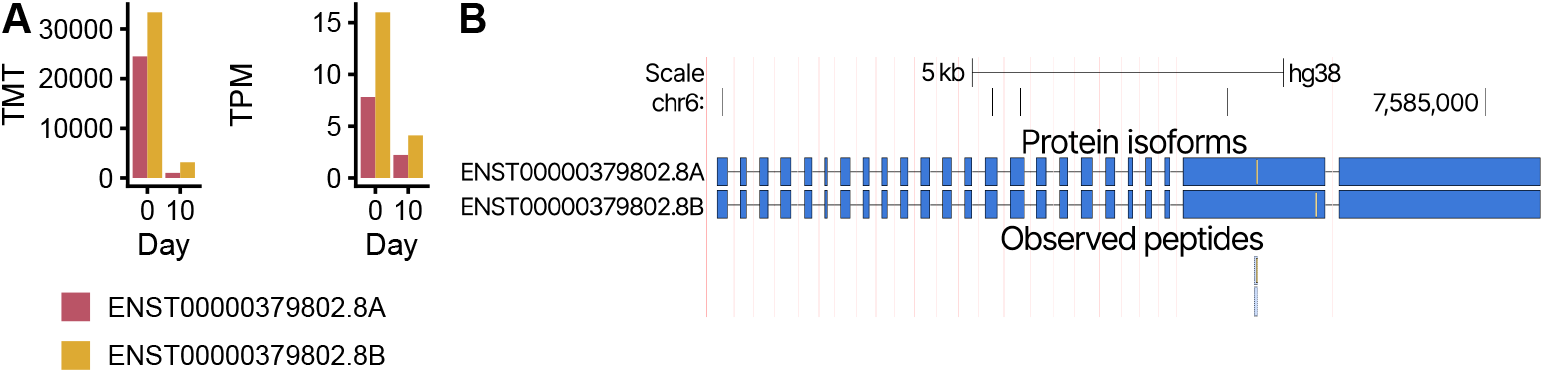
Sample-specific haplotype-resolved MS database construction enables the investigation of proteomic complexity in arbitrary models. **A.** Transcript per Million (TPM) lrRNA-seq quantifications (left) and TMT reporter ion intensities (right) for both haplotypes of the ENST00000379802.8 protein isoform on the *DSP* gene on days zero and then of the iPSC-osteoblast differentiation. **B**. Browser track highlighting the confirmed presence of both haplotypes of the ENST00000379802.8 transcript through unique peptides (shared peptides not shown).

Overall, this case study shows that our method can be used not only to describe and investigate the proteome of single samples, but also to point to differences between samples. More broadly, our method could be useful for multiple biological applications, such as the quantification of protein levels contributed by *A1AT* M-haplotypes [38].

## Discussion

Our work presents, to the best of our knowledge, the first pipeline that creates and enables the searching of sample-specific haplotype-resolved proteomes from sample-matched lrRNA-seq and MS data. We implement this framework as a Snakemake workflow that integrates existing methods where possible. We show that variant phasing for proteogenomics purposes can be performed effectively from lrRNA-seq data, especially in areas with sufficient coverage. We also characterized the theoretical and observed proteome complexity in a WTC11 cell line sample. The majority of proteome complexity was still observed to be driven by homozygous genetic variation and heterozygous genetic variation for which phasing was not required. Nevertheless, our results point toward the increased future importance of haplotype-resolved proteomes, especially as sequencing as well as MS depth and accuracy continue to improve. Finally, we highlighted the applicability of investigating sample-specific proteomes of realistic biological systems in the ten-day differentiation from iPSCs to osteoblasts.

Consistent with previous work, genetic variation contributed more to measurable proteome complexity relative to alternative splicing, both for the observed and the theoretical proteome [15]. Interestingly, the number of protein isoforms inheriting two or more heterozygous variants was relatively small, although this was likely due to the low depth of our lrRNA-seq data. There was limited linkage between genetic variation and alternative splicing, with only a handful of protein isoforms presenting both types of variation in the theoretical proteome, and none of them being identified during the search. These results suggest that the processes of genetic variation and alternative splicing may be largely independent, given that we do not find an increased proportion of alternatively spliced isoforms with variants, relative to all alternatively spliced isoforms. It should be noted that we did not attempt to identify haplotype-specific alternative splicing, as some methods have recently incorporated, which may impact this aspect [39]. Another point of note is the total number of identified variants, either via direct peptide evidence or via linkage. Interestingly, we found that the number of identified indels was much larger for the linked variants than the direct ones, both in absolute numbers and proportionally. To the best of our knowledge, this feature is caused by the fact that the direct identification of an indel requires coverage of the complete indel by a peptide, whereas linkage requires coverage of only a single single nucleotide variants (SNVs) on the same haplotype.

Our work has some limitations. First, we focused on PacBio Iso-Seq lrRNA-seq data from a WTC11 cell line in our work, as well as PacBio Iso-Seq on the iPSC-osteoblast differentiation. In addition, our benchmarks were also performed on PacBio Iso-Seq data. As such, some of our conclusions may be specific to the Iso-Seq technology or one of the two systems we investigated. Further, some of our statements regarding proteome complexity, while largely concordant with previous work, may also be a function of the lower sequencing depth of our data.

The phasing benchmark we performed was based exclusively on GIAB data. While this is standard, it is possible that some of the methods we benchmarked for variant phasing were implicitly tuned to GIAB data during development, due to its frequent usage as a development dataset, potentially skewing some of our results. Our workflow currently does not account for altered start codons. However, the number of these cases was extremely small in our experiments, occurring at frequencies comparable to altered stop codons (*<* 0.1%), but they may be more relevant in other biological contexts. Lastly, inflating the proteome size by adding potentially spurious protein isoforms may also lead to additional issues with FDR calibration [40].

A final limitation of our current pipeline concerns the mass-spectrometry level detectability of sample-specific variant and haplotype sequences. While our workflow generates and searches haplotype-resolved protein databases, we did not systematically assess whether all sample-specific variants produce peptides that are theoretically observable and confidently identifiable by bottom-up proteomics. Previous work has examined the challenges of matching peptides containing multiple variants to MS spectra and the associated effects on discoverability and identification reliability in the context of population-based haplotype databases [21, 22]. Future work could extend this analysis to sample-matched, haplotype-resolved proteomes.

There are several potential avenues for future work. First, variants phased with lrRNA-seq are now also increasingly used for other tasks, such as the quantification of ASE [29]. While our benchmark demonstrated that phasing variants with lrRNA-seq can be done accurately, this is less clear for downstream tasks such as ASE and should thus be further investigated. Our work primarily highlights the usage of haplotype-resolved proteomes for improved isoform characterization. It would also be highly desirable to investigate the implications of haplotype-resolved transcriptomes and proteomes for other tasks, such as neoantigen prediction and validation. A last and perhaps particularly interesting route for future investigation is further exploring protein (group) algorithms in the context of haplotype-resolved proteomes. Given the fact that haplotype-resolved proteomes produce a much larger number of shared peptides than typical searches, benchmarks should likely evaluate them separately [41].

## Conclusion

We have introduced a sample-specific workflow that generates and performs MS search on haplotype-resolved proteomes, based on sample-matched MS and lrRNA-seq input data. Our results show that such proteomes can be generated in practice and can support protein isoform identification at allele-specific resolution. Our work also reveals that lrRNA-seq data can not only be used to call variants accurately but also to phase them [42].

We envision this framework as an extensible foundation for lrRNA-seq–based proteogenomics. By emphasizing modularity and integration with existing tools, our workflow is well-suited for adaptation as sequencing and mass spectrometry technologies continue to advance. In this way, we believe our pipeline may be helpful both for basic research in proteogenomics as well as applications.

## Methods

### Datasets

#### Miscellaneous

##### Haplotype-resolved quantification

For haplotype-resolved quantification (Fig. 4A), haplotype-resolved transcriptomes were generated using g2gtools 3.1.0, followed by quantification of the lrRNA-seq data using lr-kallisto 0.51.1 [43].

#### WTC11 cell line

##### PacBio lrRNA-seq data

PacBio lrRNA-seq data (i.e., Iso-Seq) on the WTC11 cell line was downloaded from NCBI (SRR18130587) [42] (see Data Availability). Further experimental details are described in [42].

##### Mass spectrometry data

Bottom-up, trypsin-digested mass-spectrometry data on the WTC11 cell line was acquired directly from Bollon et al. [44] as RAW files and subsequently converted to mzML for database searching using ProteoWizard `msconvert` using the `--zlib --32 --filter “peakPicking true 1-”` parameters [45] (see Data Availability). Further experimental details are available from Bollon et al. [44] upon request.

#### iPSC-osteoblast differentiation

##### PacBio lrRNA-seq data

PacBio lrRNA-seq data (i.e., Iso-Seq) data on the iPSC-osteoblast differentiation cell line were obtained directly from the authors [36].

##### Mass spectrometry data

TMT MS data on the iPSC-osteoblast differentiation cell line were obtained directly from the authors [36].

#### Genome in a Bottle

##### lrRNA-seq data

uBAM files were downloaded from NCBI for the HG002 and HG005 Iso-Seq samples (see Data Availability).

### References

#### Reference genome

We used the `GCA_000001405.15_GRCh38_no_alt_analysis_set.fna` hg38 genome as the reference genome throughout all of our analyses.

#### Reference transcriptome

The primary assembly of GENCODE V45 was used as our reference transcriptome throughout this work [46].

#### Contaminants

We used the common Repository of Adventitious Proteins, cRAP, as contaminants in all mass-spectrometry database searches across all of our experiments.

#### UniProt

The UP000005640 human reference proteome, including Swiss-Prot and TrEMBL proteins, as well as Swiss-Prot isoforms, was used as a reference proteome, in addition to the GENCODE V45-derived proteome, in our database searches [47].

### Reproducibility

All products, including figures and tables, created and used within this paper are fully reproducible via a Snakemake pipeline available on our [Github repository](https://github.com/dnwissel/lr-haplotype-proteomics).

Experiments were run on AMD EPYC 7742 64-Core Processor CPUs under Ubuntu 22.04, using Snake-make 8.2.3 and SingularityCE 3.11.4. All other project-relevant software was run using mamba environments within Singularity containers via Snakemake. All software versions are pinned within their respective conda environments within the Snakemake pipeline and additionally logged on Zenodo (see Data Availability) [48].

### Additional pipeline details Alignment

Our pipeline begins with quality control and alignment. In particular, we first remove sequencing adapters and trim polyA tails from the lrRNA-seq data using `lima` and `isoseq refine`. Following this, we align the lrRNA-seq data to the genome using `minimap2` [49]. lima and isoseq refine were run using their default parameters. minimap2 was run using `minimap2 -ax splice:hq -uf`, as recommended by the manual for PB Iso-Seq data.

### Variant calling

We call variants based on genome alignments using `Clair3-RNA` (v0.1.0, CALLER version 0.0.2) [50], followed by appropriate filtering of the called variants using R code. Afterward, we phase the variants using WhatsHap [24, 25]. Following this, we postprocess the phased variants by removing any heterozygous variants belonging to phase blocks that do not stretch across the full CDS region in which they are located. In addition, we arbitrarily assign phase blocks to variants overlapping only CDS regions that contain solely this variant, since phasing is arbitrary in these cases.

For variant calling with Clair3-RNA, we kept both indels and SNVs. In addition, we required a minimum quality of called variants of zero, a minimum allele frequency for called SNVs of 0.08, minimum allele frequency for called indels of 0.15, a minimum coverage of three for all called variants, and a minimum mapping quality of five for an alignment to be considered.

Post-variant-calling, we filtered all variants not fulfilling a minimum GQ of five, in addition to removing multi-allelic variants.

Variant phasing using WhatsHap was performed using default parameters, except that we turned on `--distrust-genotypes`, as it has been shown to improve performance in some scenarios [26].

### Isoform discovery

Using the genome-aligned lrRNA-seq reads, along with the reference transcriptome, we perform isoform discovery using `Bambu` [51]. On the Bambu-derived transcriptome, we call open-reading frames for all novel isoforms using `orfanage` [52]. We then post-process the resulting novel and reference isoforms by filtering for valid full CDS regions and a minimum number of nucleotides.

We ran Bambu with default parameters, letting it choose a novel discovery rate, which the method set to 0.067 for our WTC11 cell line sample.

For the reference GENCODE annotation, we standardized the input GTF using `gffread` [53], adjusting stop codons (`--adj-stop`) and filtering for complete coding sequences (`-J`). Following this, we filtered the resulting GENCODE coding transcripts for a minimum CDS length of 100 nucleotides.

For ORF calling, we inputted the union of all resulting filtered GENCODE coding transcripts and the novel Bambu isoforms, calling ORFs only for Bambu isoforms. orfanage was run with default parameters, using the filtered GENCODE coding transcripts as a reference and using the `BEST` mode. Following ORF calling, isoforms were filtered for coding sequences of a minimum length of 100 nucleotides.

### Mass spectrometry database creation

To create a personalized proteome, based on our phased variants and partially novel transcriptome, we use `haplosaurus` [6]. In particular, we input a reference genome, the Bambu-derived transcriptome and the phased variants to generate protein haplotypes. We then post-process the Haplosaurus output by first refiltering the resulting protein isoforms for complete CDS region containment and a minimum codon length, since, for example, changed stop codons may have led to the truncation of some protein isoforms. Afterward, we generate unique IDs for each haplotype-resolved protein, appending A and B to the protein ID to signify the first and second haplotypes per protein isoforms. Then, we generate a sequence-unique set of protein isoforms, tracking which protein IDs result in the same protein sequence. Then, we concatenate contaminants and generate decoys for use in database searching using `DecoyPYrat` [54], to finalize the protein database.

Haplotype labels (A, B) are assigned per CDS, aiming to keep haplotypes together that inherit the same genotype of a particular heterozygous variant. There are some cases where the same heterozygous variant is assigned different haplotype labels, as haplotype labels are assigned per isoform, without looking back at all previously created labels. We’ve found these cases to be rarer than not and not to deter post-search visualization and annotation.

DecoyPYrat was run using pypgatk 0.0.24 for trypsin, allowing a maximum of two missed cleaves, with a minimum peptide length of five and a maximum peptide length of 100, for a maximum of 100 iterations for decoy generation.

### Mass spectrometry search

We then perform a database search using `sage` 0.14.7 [55], inputting our generated MS database and the sample MS data. Database searches were performed separately for (i) the sample-specific haplotype-resolved proteome, (ii) the GENCODE V45 reference proteome, and (iii) the UniProt reference proteome (UP000005640). All searches used identical search parameters unless otherwise specified below.

For the WTC11 cell line searches, sage was configured with the following parameters: trypsin digestion with a maximum of two missed cleavages; peptide length range of 5-100 amino acids; peptide mass range of 500-5000 Da; precursor mass tolerance of 10 ppm; fragment mass tolerance of 10 ppm; precursor charge states 2-4; static modification of carbamidomethylation on cysteine; variable modification of oxidation on methionine with a maximum of two variable modifications per peptide; minimum of 15 peaks and maximum of 500 peaks per spectrum and a minimum of 4 matched peaks for identification.

For the iPSC-osteoblast differentiation TMT-labeled searches, sage was configured with TMT16-specific parameters: trypsin digestion with a maximum of two missed cleavages; peptide length range of 5-100 amino acids; peptide mass range of 500-5000 Da; precursor mass tolerance of *±*10 ppm; fragment mass tolerance of *±*0.1 Da; precursor charge states 2-4; static modifications of TMT16 on peptide N-terminus and lysine and carbamidomethylation on cysteine; variable modifications of oxidation on methionine and phosphorylation on serine, threonine, and tyrosine with a maximum of two variable modifications per peptide; minimum of 15 peaks and maximum of 500 peaks per spectrum; minimum of 4 matched peaks for identification; retention time prediction enabled; and TMT16 quantification with level 3 settings.

Based on the search results, we perform protein inference using a standard greedy set cover approach [56] that we implement using `RcppGreedySetCover` [57] in R. In addition, we create BAM files that enable the creation of browser tracks for easy visualization of the search results by extracting genome-space sequences of both protein isoforms and observed peptides in R. Lastly, we perform post-search annotation in R, aggregating information from isoform discovery, variant calling, and protein haplotype generation to provide detailed information on if and how each protein isoform differs from the set of reference protein isoforms.

Full sage configuration files are available on Github and Zenodo for reproducibility (see Code Availability).

Search using sage was run using identical parameters as the decoy generation, where applicable. Full sage configuration files are available on Github and Zenodo for reproducibility (see Code Availability).

All post-search annotation was performed using an FDR of 0.01, using the q-value estimates produced by sage. Browser tracks for protein isoforms and matching peptides are created via custom matching in R to back-translate protein sequences to transcriptomic sequences, accounting only for non-silent variation. The resulting FASTA files were then assigned to a haplotype (homozygous, first haplotype, or second haplotype). This is trivial for protein isoforms. For peptides, we assigned them to a particular haplotype if they only matched one haplotype of one protein isoform and assigned them to be homozygous in all other cases. Protein isoform sequences were aligned to the genome using minimap2, using standard parameters for the spliced alignment of lrRNA-seq while peptide sequences were aligned using STAR [58], due to their short length.

The annotation file contains the following information per haplotype-resolved protein, constructed primarily out of Haplosaurus and SQANTI protein output files, and aggregated via R code:

1. Protein splice category as determined by SQANTI protein [4].
2. Number of contained non-silent genetic variation, broken down by SNV/indel and homozygous/heterozygous.
3. List of all contained non-silent genetic variation.
4. Diff string between the observed protein isoform and the reference without any genetic variation.
5. Flags for several protein effects, including indel, changed stop codon, frameshift, and resolved frameshift.

### Benchmark structure

We performed a phasing-only benchmark, in which we re-phased GIAB data for the HG002, and HG005 GIAB cell lines and estimated phasing accuracy- and completeness, relative to a ground truth of ancestry-derived phasing. Importantly, this benchmark evaluates only phasing, as we reuse the true unphased genetic variation as an input.

### Preprocessing

To prepare the GIAB datasets for variant calling and phasing, we preprocessed them in the following way.

#### PB Iso-Seq

We performed adapter removal and polyA tail trimming using lima and isoseq refine. Following this, data was aligned to the reference genome using standard settings for Iso-Seq data (`minimap2 -ax splice:hq -uf`).

#### Phasing-only benchmark

In our phasing-only benchmark, we considered datasets from GIAB, HG002, and HG005, evaluating PB Iso-Seq for each.

As performance metrics to evaluate phasing, we considered both accuracy and phasing completeness. We measured accuracy via the switch error rate as computed by WhatsHap `compare` and phasing completeness using both the total and relative number of phased variants as well as the total and relative number of fully phased regions of interest, which we consider to be CDS-, transcript- and gene-level.

In addition, similar to previous work on variant calling with lrRNA-seq, we stratify results by the required minimum coverage a considered variant has to have in the lrRNA-seq data to be included in the benchmark [42]. We considered minimum input coverages of 5, 10, 25, and 100. Furthermore, we stratified results by whether we phased only SNVs or SNVs and indels jointly, as this has been shown to impact performance for variant phasing [24].

#### Phasing methods

All methods were run with default parameters unless otherwise noted.

#### WhatsHap

We turned on the `--distrust-genotypes` option, as it has been shown to increase performance in some cases [26]. WhatsHap was ran using version 2.2.

#### HapCut2

`extractHAIRS` was run with the `--ont` flag enabled, as this is recommended in the manual for long-read data. HapCut2 was ran using version 1.3.4.

#### Margin

Margin was run using PEPPER-DeepVariant r0.8.

## Acknowledgements

The authors thank Madison M. Mehlferber and Elizabeth Tseng for helpful feedback and discussions throughout the project. The authors thank Saikat Bandyopadhyay for feedback on running a proof-of-concept iteration of the pipeline on Singularity. We would also like to acknowledge Mayank Murali for general feedback on an earlier iteration of the pipeline, Marina Panteli for exploring early possibilities for incorporating genetic variant information into long-read-derived proteomes, and Roger Volden for general feedback on the project.

## Declarations

## Ethics approval and consent to participate

This study used only previously sequenced, publically available datasets and was thus exempt from ethics approval.

### Consent for publication

Not applicable.

### Data availability

#### Raw data

Raw lrRNA-seq data for the WTC11 cell line is available from [NCBI](https://www.ncbi.nlm.nih.gov/sra/SRR18130587) (SRR18130587). Raw MS data for the WTC11 cell line are available from Bollon et al. [44] upon request.

Raw lrRNA-seq for the GIAB datasets is available as follows:

1. HG002 PB Iso-Seq: [NCBI](https://ftp-trace.ncbi.nlm.nih.gov/ReferenceSamples/giab/data_RNAseq/AshkenazimTrio/HG002_NA24385_son/Baylor_PacBio/reads/m64139_220127_180020.hifi_reads.bam).
2. HG005 PB Iso-Seq: [NCBI](https://ftp-trace.ncbi.nlm.nih.gov/ReferenceSamples/giab/data_RNAseq/ChineseTrio/HG005_NA24631_son/Baylor_PacBio/reads/m64139_220129_000012.hifi_reads.bam).

#### Reference data

Reference data is downloaded as part of our Snakemake pipeline (see Code availability). In addition, the GENCODE V45 primary assembly is available from [GENCODE](https://www.gencodegenes.org/human/release_45.html), the hg38 reference genome from [NCBI](https://ftp.ncbi.nlm.nih.gov/genomes/all/GCA/000/001/405/GCA_000001405.15_GRCh38/seqs_for_alignment_pipelines.ucsc_ids/GCA_000001405.15_GRCh38_no_alt_analysis_set.fna.gz), the cRAP contaminants from [GPM](http://thegpm.org/crap/), and the [UniProt](https://www.uniprot.org/proteomes/UP000005640) human reference proteome from UniProt.

In addition, GIAB phased genetic variation files are available from NCBI for HG002 and HG005.

#### Produced result data

Result data is cached and publically available on Zenodo [48].

### Materials availability

Not applicable.

### Code availability

All code necessary to reproduce the results in our paper is available on our [Github repository](https://github.com/dnwissel/lr-haplotype-proteomics). All code has, in addition, been cached on [Zenodo](https://doi.org/10.5281/zenodo.17674329) Zenodo, where it is also freely accessible [48].

Code to apply the pipeline on user-provided samples is available on a separate [Github repository](https://github.com/dnwissel/lr-haplotype-proteomics-pipe).

### Author contribution

Study conception: GMS, MDR. Study design: DW, GMS, MDR. Benchmark design: DW. Software implementation: DW. Analysis and interpretation of results: DW, GMS, MDR. Draft manuscript preparation: DW, GMS, MDR. All authors reviewed the results and approved the final version of the manuscript.

---

**Fig. S1.**
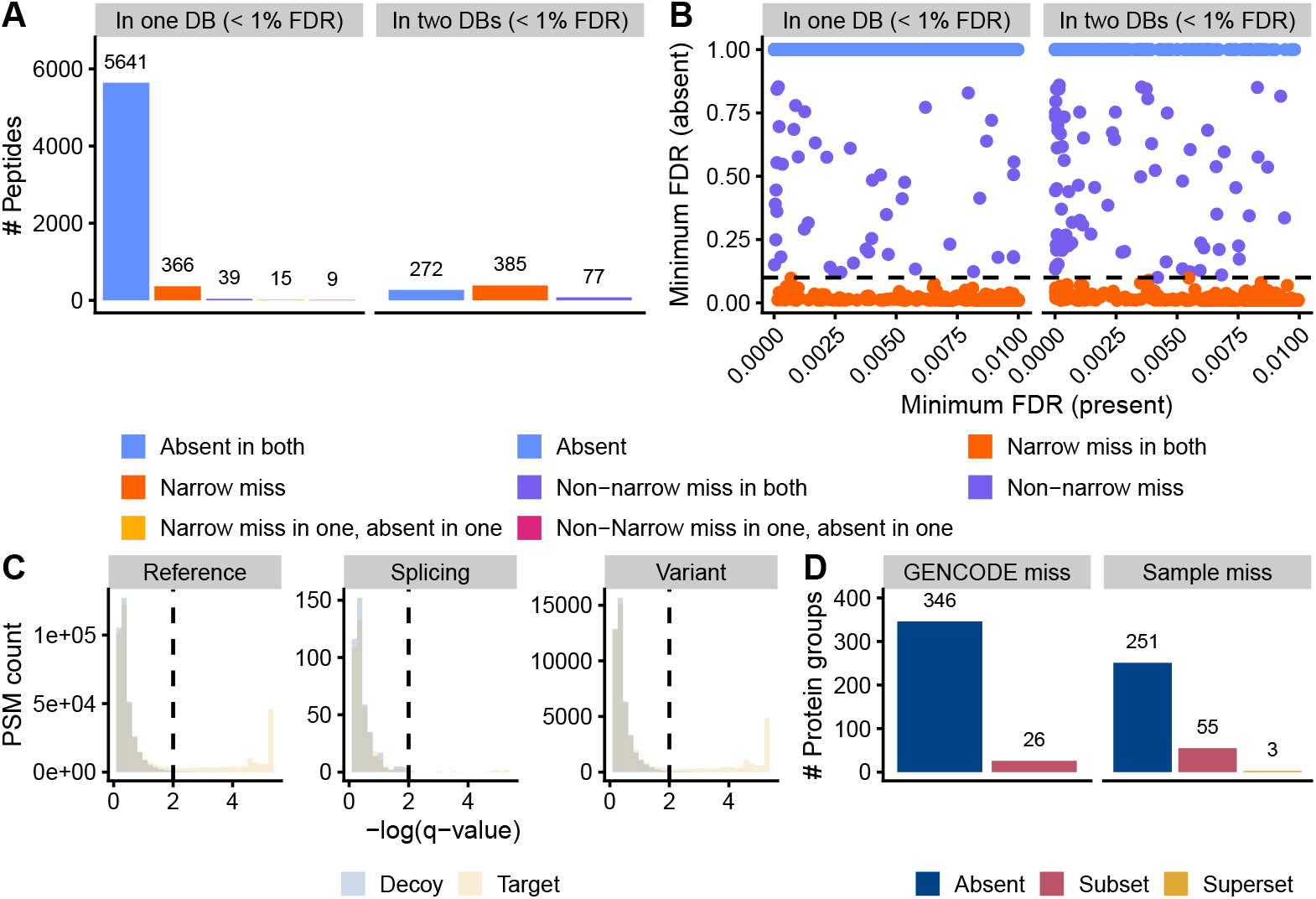
Reasons for peptide differences between different databases, calibration of decoy-target FDRs in our sample-specific database, and protein group differences between our sample-specific, GENCODE-derived database and the GENCODE reference. **A.** Discrete classification, taking into account whether the peptide in question behaved similarly in databases in which it was not detected. **B**. Quantitative classification, using the minimum q-value for databases in which the peptide was detected (if the peptide was detected in two databases) or peptides in which the peptide was not detected (if the peptide was detected in a single database). Absent in both indicates that the peptide did not have any PSMs in the missing database(s), independent of q-value. Narrow miss in both and narrow miss indicate that the peptide had at least one PSM with a q-value *q <* 0.1 in the missing database(s). Non-narrow miss in both and non-narrow miss indicate that the peptide had at least one PSM with a q-value *q >* 0.1 in the missing database(s). Narrow-miss in one, absent in one, and non-narrow miss in one, absent in one each indicate that the peptide in question was absent in one of the missing databases and had a narrow or non-narrow miss in the other (definitions as before). **C**. Calibration of estimated q-values in the target-decoy framework on the WTC11 cell line, stratified by the source of the peptide. Only peptides mapping to a single protein isoform were considered. Peptides were assigned to the reference peptide group unless the protein isoforms they mapped to were alternatively spliced (in which case they were assigned as splicing peptides) or had at least one non-silent variant (in which case they were assigned as variant peptides). Peptides that could be counted as either variant or splicing peptides were counted as splicing peptides. The dotted line denotes *q* = 0.01. **D**. Reasons for protein group differences between the GENCODE reference proteome and the WTC11 sample-specific proteome, stratified by the database in which protein groups were missing. Absent denotes that at least one protein of the protein group was absent from all inferred protein groups for the other database. Subset indicates that the protein group in question was a subset of a protein group in the other database. Superset indicates that the protein group in question was a superset of a protein group in the other database.

